# Meta-learning leading to homeostatic plasticity stabilizes synaptic weights together with predictable activity levels

**DOI:** 10.64898/2026.06.16.732795

**Authors:** Florentin Wörgötter, Konstantin Möller, Minija Tamosiunaite

**Author notes:** These authors have contributed equally to the work (Florentin Wörgötter, Konstantin Möller).

## Abstract

Stabilizing synaptic plasticity together with the neuron’s activity has remained a central challenge in theoretical neuroscience since the introduction of Hebbian learning principles. Classical Hebbian learning rules typically lead to unbounded synaptic growth, motivating the development of stabilization mechanisms such as normalization methods, BCM-type learning, synaptic scaling and others. While these approaches can prevent divergence, they can also exhibit different limitations e.g. resulting in too-sparse synaptic configurations or leading to poor scalability with increasing network size. A recently introduced meta-plasticity mechanism, termed annealed linear learning (ALL), dynamically reduces the learning rate as neuronal output increases, thereby preserving stable and interpretable fixed-point behavior of the output. However, the original formulation leads to an irreversible decay of the learning rate, preventing adaptation to changing environmental conditions. To address this, in the present study, we balance learning rate reduction at large outputs with recovery at small outputs and in addition introduce forgetting that gradually reduces synaptic weights. These extensions allow the system to discard outdated representations and adapt to novel input conditions. Analytical investigations demonstrate that the favorable output fixed-point properties of the original ALL framework are preserved under the extended rule. Furthermore, simulations with an artificial agent show that the proposed mechanism enables robust and fast re-learning and adaptation in changing environments.

## Introduction

Stabilizing synaptic plasticity together with neuronal activity has been an issue in the theoretical neuroscience ever since mathematical plasticity rules had been designed starting from Donald Hebb’s famous conjecture [1] often abbreviated by “what fires together wires together”.

Conventional Hebbian learning leads to unbounded (divergent) weight growth and an unrealistic increase of the neuronal output. Many stabilization methods and/or augmentations of the original Hebbian learning rule have been suggested to prevent this, for example Oja’s rule [2], the Bienenstock, Cooper, Munro rule (BCM, [3]), subtractive normalization methods [4], and several more.

As an alternative, researchers have discussed spike timing dependent plasticity [5–7], which leads to long term potentiation when post-synaptic follows pre-synaptic activation and to long term depression when the order of signals is reversed.

More recently, learning rules had been introduced which combine Hebbian weight growth with a homeostatic, balancing term, called synaptic scaling [8–10], for achieving convergence to a target activity [11].

All of these rules can stabilize synaptic weights but they often show certain disadvantages. For example, subtractive normalization can lead to overly strong competitive effects resulting in the extreme case in one synapse with a high value whereas the others are (near) zero.

BCM displays a fix point characteristic of the final activity, which is - as such - an advantage but in a multi-input situation fix points at the output do not reflect the strength of the inputs and their combinations at all [12]. Furthermore, convergence of BCM will slow down exponentially when more and more synapses are modeled [13], which can render BCM practically useless in larger networks.

Synaptic scaling had been an attractive idea supported by experiments as it can stabilize synapses and output at the same time. In a nutshell is says that neurons “love to fire not too much and not too little” trying to adjust synapses such that their firing will converge to some target activity *v*_*T*_ [8–10]. Weight change 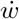 following input *u* based on plasticity combined with scaling usually is implemented with: 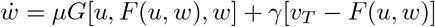, where *G* abbreviates a plasticity rule (such as Hebb or BCM) and *F* is a general neuron model where the neuron’s activity *v* is just *v* = *F* (*u, w*). The first term, thus, represents plasticity and the second synaptic scaling. Constants *γ* ≪ *µ* ≪ 1 are the rate-factors of both processes. It could, however, be shown that stability requires introducing an additional weight dependence into the scaling term as: 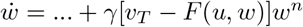, where *n* ≥ 2 is needed without which many conventional plasticity rules *G* such as BCM remain unstable [11]. Alas, for this there is so far no experimental support. Furthermore, we have observed (see e.g. [12, 14]) that in network implementations of the combined plasticity+scaling rule many times *v*_*T*_ ≤ 0 is needed to obtain stability. Naturally, the concept calling *v*_*T*_ “target activity” is thwarted this way.

It would, thus, be desirable to arrive at a learning rule with predictable output fix point characteristic that reflects the input structure and remains in a range compatible with the experimentally supported notion of a target output activity.

To achieve this, we had in a recent study introduced a meta-plasticity mechanism (“annealed linear learning”: ALL-rule) that adds to a learning rule a mechanism for reducing its rate factor *µ* as soon as the neuron’s output is large [12]. This relates to known effects where LTP is smaller for large synapses as compared to small synapses [15]. As a disadvantage, this rule leads to the final situation that the learning rate drops to zero and cannot be recovered, which prevents a system from re-learning for example in a situation where the input situation changes, due to a change in the environment.

In this case, you should forget what you have learned and rather learn about the new situation. To achieve this we have in the current study introduced a forgetting mechanisms that reduces weights. Furthermore, we can recover the learning rate allowing for the learning of a changed (new) situation. We will analyze this rule analytically demonstrating that this system generates predictable activity fix points leading to an effect akin to synaptic scaling [8–10] and show, in addition, that an artificial agent can now adapt to novel situations.

## Materials and methods

### Learning Rule

The system we are investigating relies on, and substantially modifies, the so-called *annealed linear learning (ALL)* rule [12]. In the following we calculate the properties of our system for two inputs *u*_1_ and *u*_2_, which is the relevant situation for the simulated closed loop behavioral system discussed later. The case for a single input can be determined in a straight-forward way by setting one of them to zero. Hence, we define the output of the neuron *v* as:

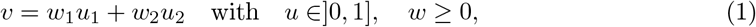

where *w* are the synaptic weights and *u* the inputs. When we omit indices then an equation will hold for both inputs.

The conventional Hebb-rule reads as *dw/dt* = *µuv*, with *µ <<* 1 the learning rate. To define the ALL-rule we use, instead, a Heaviside function for the *v*-term, defining a linear Hebb-rule as:

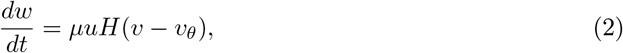

where *H* will be 1 as soon as *v > v*_*θ*_ and zero else. For simplicity we set *v*_*θ*_ = 0 and get

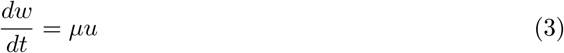

The reduction of the conventional Hebb-rule to the here-used linear variant has been in-depth analyzed in [12]. It is justified physiologically by the fact that especially at dendritic spines *Ca*^++^ influx through NMDA channels, which determine LTP, have an all-or-none effect on plasticity [16, 17]. As a consequence, it appears that every post-synaptic back-propagating spike or dendritic spike will be enough to lead to substantial *Ca*^++^ influx to trigger plasticity (at a spine). This argues for a sharp – maybe sigmoidal – transition of the post-synaptic learning influence, where the use of the Heaviside function would represent a limit case. In the Discussion section we will in more depth consider the physiological background for this.

Then we add a forgetting term −*fw* and and a time constant *τ*_*w*_ and change this to:

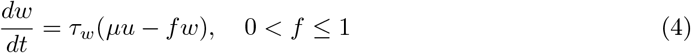

Finally we define a function that implements meta-learning by changing the learning rate *µ* using two opposing processes. The learning rate will get reduced for large output *v* using a process similar to simulated annealing as introduced in [12]. However, when *v* drops too much a recovery term will bring the learniong rate up again to allow for learning to start again in case of changing inputs. For this we define (indices “a,r” for annealing,recovery, respectively):

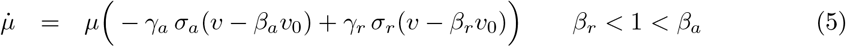

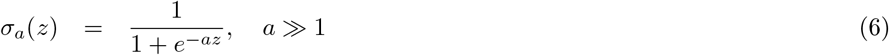

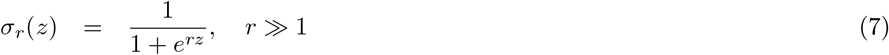

with 0 *< γ*_*a,r*_ *<* 1. The sigmoidal functions *σ*_*a*_ and *σ*_*r*_ bring the learning rate rather suddenly down or up when *v* is clearly above or below *v*_0_ as determined by the two shifting parameters *β*_*a*_ and *β*_*r*_.

Figure 1 shows examples of the two combined sigmoidal functions, which influence the change of the learning rate. The left side of these curves represent the recovery process which gets stronger the more *v* approaches zero, while the right side represents reduction of the learning rate (annealing) for large *v*.

**Figure 1.**
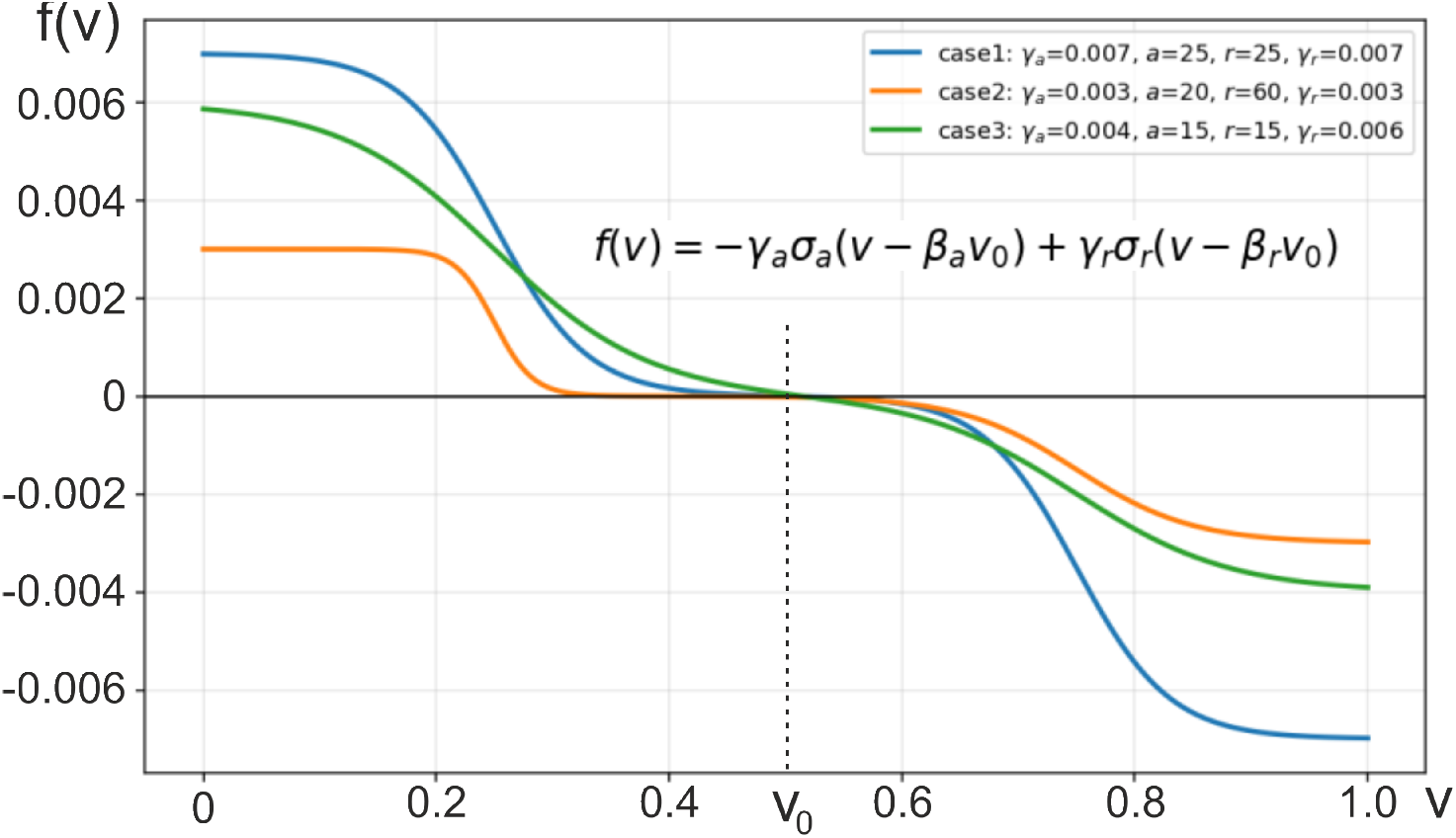
Examples of the dual sigmoids in the 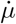 equation. Parameters: *v*_0_ = 0.5, *β*_*a*_ = 1.5, *β*_*r*_ = 0.5, all others as in the plot. For these functions we find that they approach *γ*_*r*_ for *v* = 0 and −*γ*_*a*_ for *v* = 1 (see e.g. green curve).

The complete system is, thus, given by:

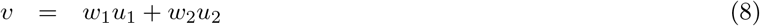

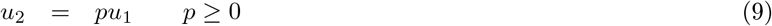

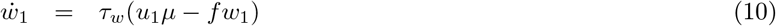

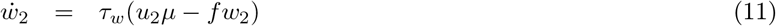

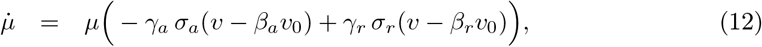

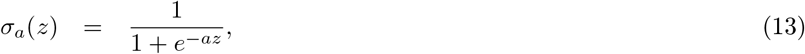

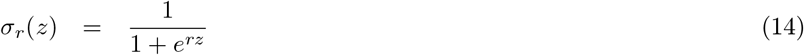

where we assume that *u*_2_ is a scaled version of *u*_1_ and *p* = 0 corresponds to the single-input case.

Note that the strength of the different terms in this rule must be balanced to allow for fast convergence, while simultaneously exhibiting plasticity. When assuring this, this rule leads to weight stabilization while it also allows re-learning in case of changes in the inputs (e.g. due to a change in the environment of an agent). This will be analyzed later by determining its fix points and with simulations (see Results).

### Experimental setup

The goal of the system is to enable agents to exhibit foraging behavior to move toward a food source in a square arena after exploring the environment. Furthermore, agents should be able to learn a new food source, when the situation has changes. The only stimuli that the agents get are the odor emanating from a food source and location information from place fields. We would like to emphasize that for these simulations we are not attempting to implement a highly realistic model of hippocampal place cells/fields [18] (see Discussion). The goal is rather to show that theoretical results obtained with 2 inputs only (see below) will still hold for larger networks with many more inputs. For this we define an (*x, y*) arena with size of numerically 1.0 × 1.0 each.

### Odor- and Location-Stimuli

The odor plume is modeled as radially emitting from the food source and decreasing in strength with increasing radial distance *ζ* from the food source with a decay that follows 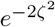. Hence at the source it takes a value of one and it spreads into the full area with a strength that decays with distance. Odor input *S*_*O*_ will, thus, take values 0 *< S*_*O*_ ≤ 1.

In addition to this single odor stimulus we are simulating a place field system to allow an agent to obtain location information. Hence, place cells get activated when the agent is inside a certain region (called “place field”) of the environment, where activation is strongest in the center of the place field creating a mapping of location to activation. Place field centers are distributed randomly for each agent via Poisson disk sampling with a fixed radius of 0.075 units, such that they cover most of the arena. We then add uniform noise *ϵ* ∈ *U* (0.05, 0.1) to the radii to get overlapping place fields with varying size. On average we obtain *>* 100 place fields this way. Note that we had also implemented place fields with variable radii and overlap, but in general result were not substantially affected by this.

To calculate the activation *S*_*L*_ (index L for “location”) of a place cell for an agent located at position (*x, y*), we measure the agent’s distance *d* from its place field center and apply a sigmoid function to obtain the activation value:

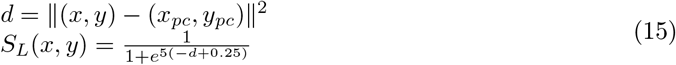

Here, (*x*_*pc*_, *y*_*pc*_) is the center location of the place field. Hence, place field inputs *S*_*L*_ will also take values 0 *< S*_*L*_ ≤ 1.

### System architecture

#### Layer 1 Place cells

The overall architecture, illustrated in Fig. 2, is organized hierarchically from the bottom up. The first place cell layer is composed of several “stacks,” each containing *N* = 200 neurons. Every stack is associated with a specific 2nd layer place cell, which serves as a higher-level integrator.

**Figure 2.**
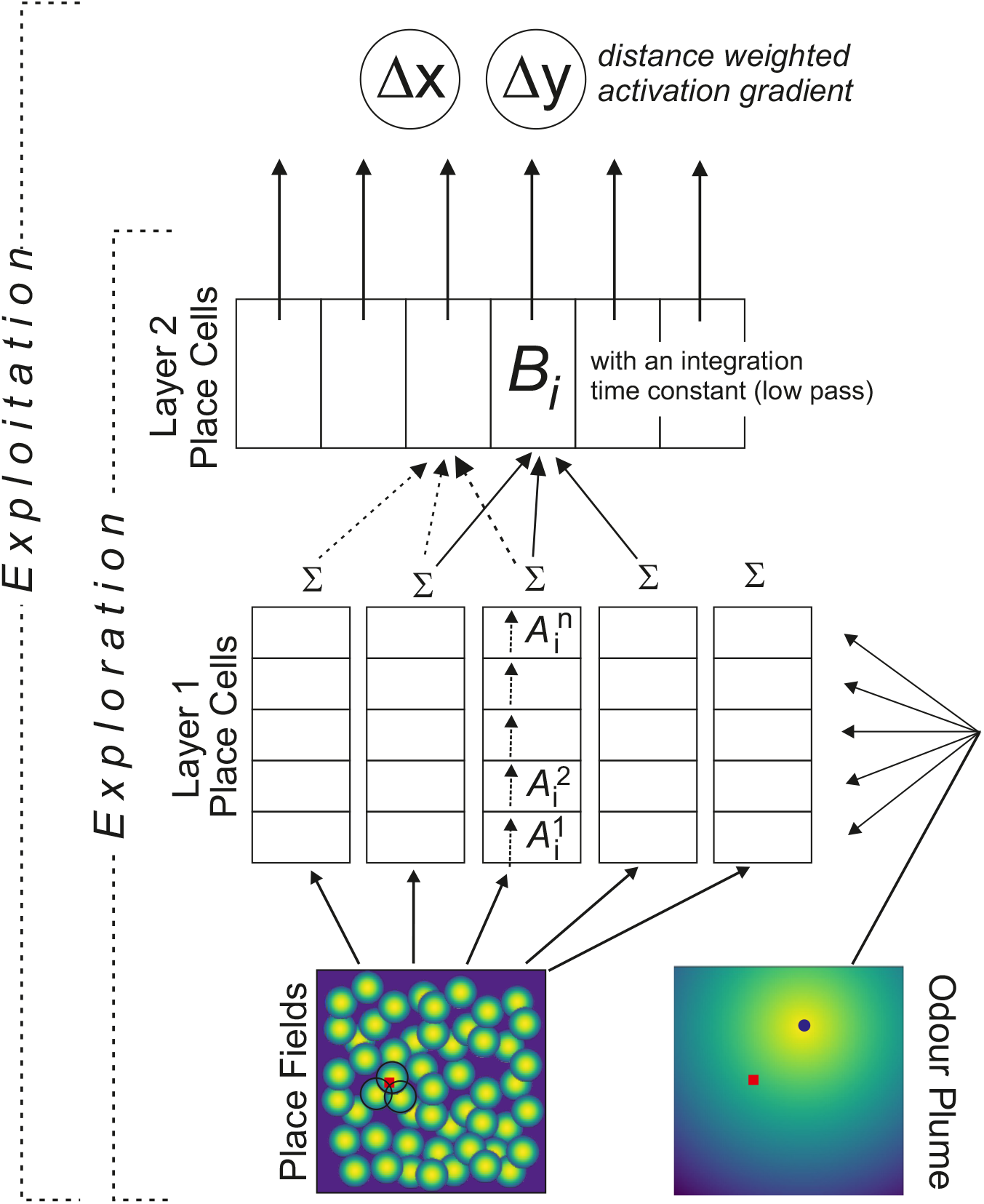
Schematic overview of the model architecture. From bottom to top: The dark blue dot in the odor panel marks the food source and the red squares the agent’s location in the environment. In this case 3 place cells will be activated (circles). The dashed vertical arrows near *A*_*i*_ indicate that any one stack of the layer 1 place cells gets input from the same place field. Stacks contain 200 neurons each. Their activity is given by Eq. 16. Place cells in the 2nd layer receive pooled inputs from nearby stacks and perform a weak temporal low pass filtering, which smoothens activities *B*_*i*_. During exploitation an activation gradient is calculated from 2nd order place cells inversely weighted by their distance from the agent. This gradient is used to determine the next movement step of the agent.

Neurons within a stack are activated by inputs that share a common place field, and, due to the odor spread, they will also receive some odor input. To achieve this we translate the originally continuous values 0 *< S*_*O*_, *S*_*L*_ ≤ 1 into probabilities of activating a subset of neurons in a given stack. For example, a stimulus strength of *S*_*L*_ = 0.4 would mean an average of 40% of the layer 1 place cells in the same stack get activated by the place field stimulus. Thus, the actual inputs *Ŝ* to a neuron in a stack are then binary: either *Ŝ* = 1 with a probability *S*_*L,O*_ or *Ŝ* = 0 vice versa.

The activation *A* for a specific place cell within as stack is then calculated as the weighted sum of its two inputs:

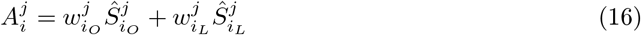

In this expression *i* denotes the index of the stack (corresponding to a specific layer 2 place cell, see next), *j* identifies the *j*-th neuron within that stack and *w* represents the synaptic weight for input *Ŝ*. As a result at any one point in time each cell in a stack can only produce 4 output values: 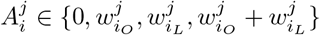. In this way we emulate in a simple manner probabilistic spiking activity and its translation into 4 possible membrane potential amplitude values.

Weights *w* change following the modified ALL rule as given by Equations 41-45. For all simulations shown below we use:

*a* = 50, *r* = 50, *γ*_*a*_ = 10^−1^, *γ*_*r*_ = 10^−2^, *β*_*a*_ = 0.8, *β*_*r*_ = 0.25, *τ*_*w*_ = 1, *f* = 10^−2^, *v*_0_ = 1.0 and units in the system are updated whenever the agent touches the place field.

#### Layer 2 Place Cells

The second layer is only used to arrive at smoother activations. Thus, we calculate an update of the activation *B*_*i*_ of the layer 2 place cells by computing a spatially weighted average of the activations *A*_*k*_ of all stacks *k* with a place field center *x*_*k*_ that is within a distance of 0.2 to the place field center of the layer 2 place cell *x*_*i*_ considered. Let *n* be the number of stacks found to obey this distance criterion given as Δ_*k*_ =| *x*_*k*_ − *x*_*i*_ |≤ 0.2, then:

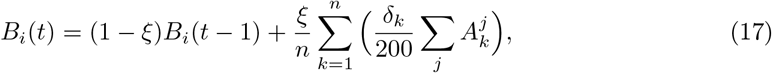

with *ξ* = 0.2 and a spatial averaging kernel given by 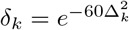. Note that each stack *k* consists of 200 neurons (see above). This combines a spatial average with a mild temporal low pass using past values of *B*_*i*_ and leads to the smoothing of the outputs from the layer 1 place cells. By this we simulate the place field of the layer 2 place cell.

### Exploration mode

To address learning and re-learning first the agents need to explore the environment. For this, the agent moves around in the arena in an exploratory manner using a direction-preserving random walk, adapted from [19]. During this the system learns.

In essence, at each timestep the agent randomly picks an angle from *ϕ* ∈ {0^◦^, 45^◦^, 90^◦^, …, 315^◦^} (0^◦^ is north with reference to the arena) and moves 0.1 units in that direction. Should the agent leave the confines of the arena, it is reset to the boundary and the heading is randomized. The probability distribution for the chosen angle approximates a Gaussian, with the center of the bell on the angle of movement from the previous timestep. The benefit of this scheme compared to e.g. random walks are the resulting more natural paths which consist of longer straight segments as in real animals. See [19] for a comparison of paths generated by this algorithm with those of real rats.

### Exploitation mode

After exploration, the agent switches into exploitation mode. To know the direction in which the agent should move, we calculate an approximate gradient (Δ^*x*^, Δ^*y*^) for each 2nd order place cell. We employ a simple scheme for this, which does not attempt to be physiologically realistic as we need exploitation only for statistical purposes (see Fig. 10).

Hence, for an agent that is located closest to a certain second order place field *c*, this is achieved in the following way. First we calculate a distance-depending weighing factor *ω*^*d*^ to make sure that place fields *i* closer to the agent exert a stronger influence on the agent than those further away:

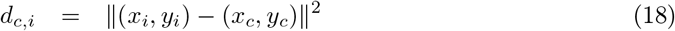

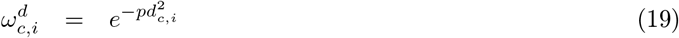

where (*x, y*) are the place field center coordinates of place fields *c* and *i*. The exponent *p* has been set to 15 empirically to render smooth gradients, but other values will also work. The weights are then normalized, such that the sum is 1. Then, we also determine the angle *θ*_*c,i*_ under which both place field centers appear as:

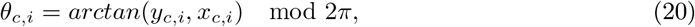

where *y*_*c,i*_, *x*_*c,i*_ are the two relevant sides of the enclosing triangle. From this we calculate the relevant x- and y-direction dependent weights that place field *i* will exert onto place field *c*. This is done simply using a cosine function on *θ*_*c,i*_ because this renders strongest impact (*cos* = 1) when place fields are aligned along the considered *x* (or *y*) direction and zero impact when they are orthogonal:

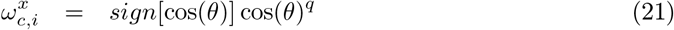

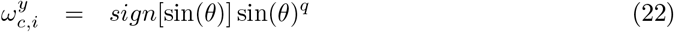

An exponent of *q* = 10 has been determined experimentally to achieve strong enough gradients, which makes it necessary to put the *sign* function in front to keep the original behavior of the cosine. Finally we get gradients Δ from this considering all *i*:

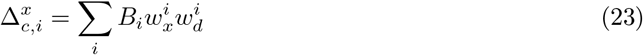

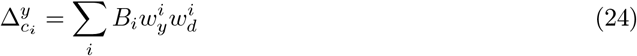

where *B*_*i*_ are the activations of place cells *i* (Eq. 17). See Figure7 B for an example of such a gradient field.

## Results

### Analytical Results – Fix Point Analysis

The main result which we will obtain in this section is that this system produces an input-independent fix point of the neuron’s output akin to the property of synaptic scaling. To show this, we consider the system given by Eqs. 39-45. For this fixpoints 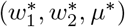 satisfy 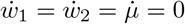. At first we note that:

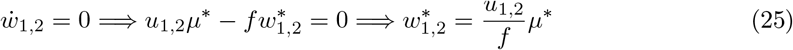

Using *u*_2_ = *pu*_1_ this implies 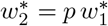.

Now insert 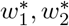 into *v* = *w*_1_*u*_1_ + *w*_2_*u*_2_:

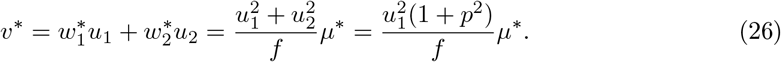

Hence

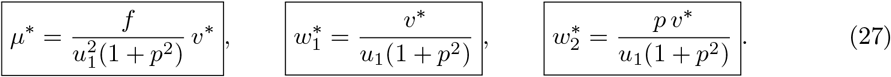

Moreover:

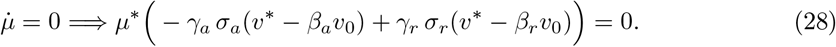

A. **Trivial fixed point:** 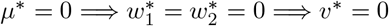. Thus, the trivial fix point is:

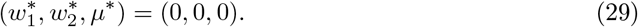
B. **Nontrivial fixed point:** *µ*^∗^≠ 0 =⇒ −*γ*_*a*_ *σ*_*a*_(*v*^∗^ − *β*_*a*_*v*_0_) + *γ*_*r*_ *σ*_*r*_(*v*^∗^ − *β*_*r*_*v*_0_) = 0.

Let *x* = *v*^∗^ − *β*_*a*_*v*_0_. Then *v*^∗^ − *β*_*r*_*v*_0_ = *x* + (*β*_*a*_ − *β*_*r*_)*v*_0_, and:

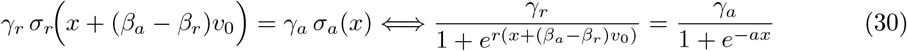

This yields:

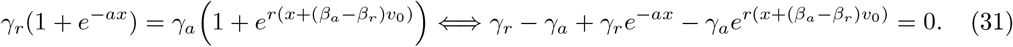

Equivalently, with *y* = *e*^*x*^ *>* 0,

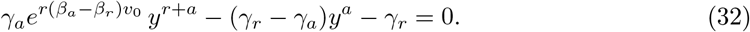

Any positive root *y*^∗^ gives

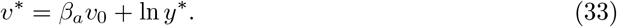

In summary:

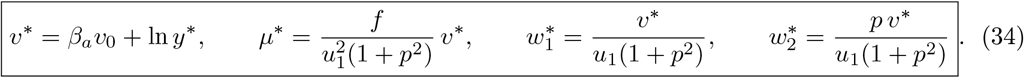

One core result here is that the activity fix point *v*^∗^ is independent of the inputs emulating a rigorous scaling principle. Furthermore, with some additional arguments (see Appendix) we find that lim_*a*→∞_ *v*^∗^ = *β*_*a*_*v*_0_. This can be seen for the special case of two identical sigmoids (*a* = *r*) considered next.

**Special case** *a* = *r*:

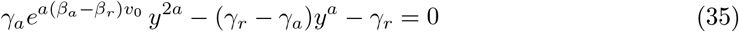

Let *z* = *y*^*a*^ *>* 0. Then

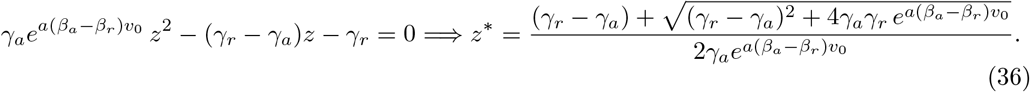

Parameters:

*a* = *r* = 45.0, *γ*_*a*_ = 10^−3^, *γ*_*r*_ = 10^−2^, *β*_*a*_ = 2.0, *β*_*r*_ = 0.25, *v*_0_ = 0.5, *τ*_*w*_ = 1.0, *f* = 10^−3^. Initial conditions were *w*_1,2_(0) = 1.0 and *µ*(0) = 10^−3^. Note that *v*^∗^ = 0.588 which is very close to the numerically calculated average of 0.571.

Hence:

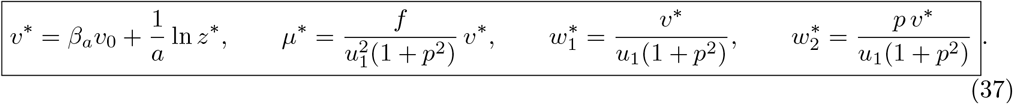

In the Appendix we show that fix points are stable. For creating Figures 3 and 4 we use — if not mentioned differently in the figure legend —the following parameters:

**Figure 3.**
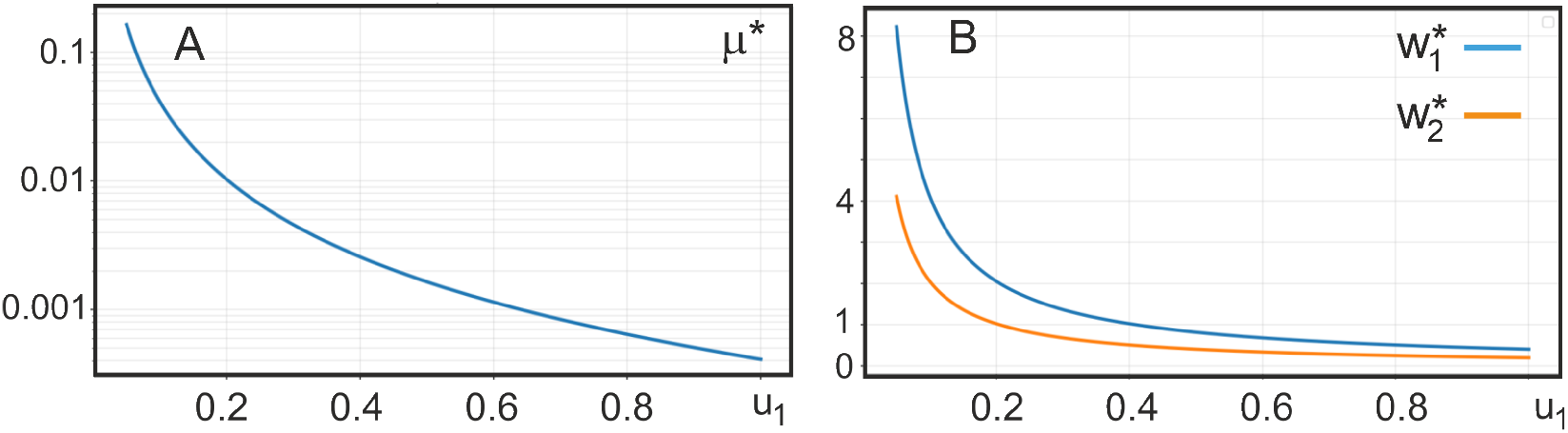
Fix points *µ*^∗^ and *w*^∗^ for the case *a* = *r* according to Eq. 37 depending on *u*_1_. For parameters see Eq. 38.

**Figure 4.**
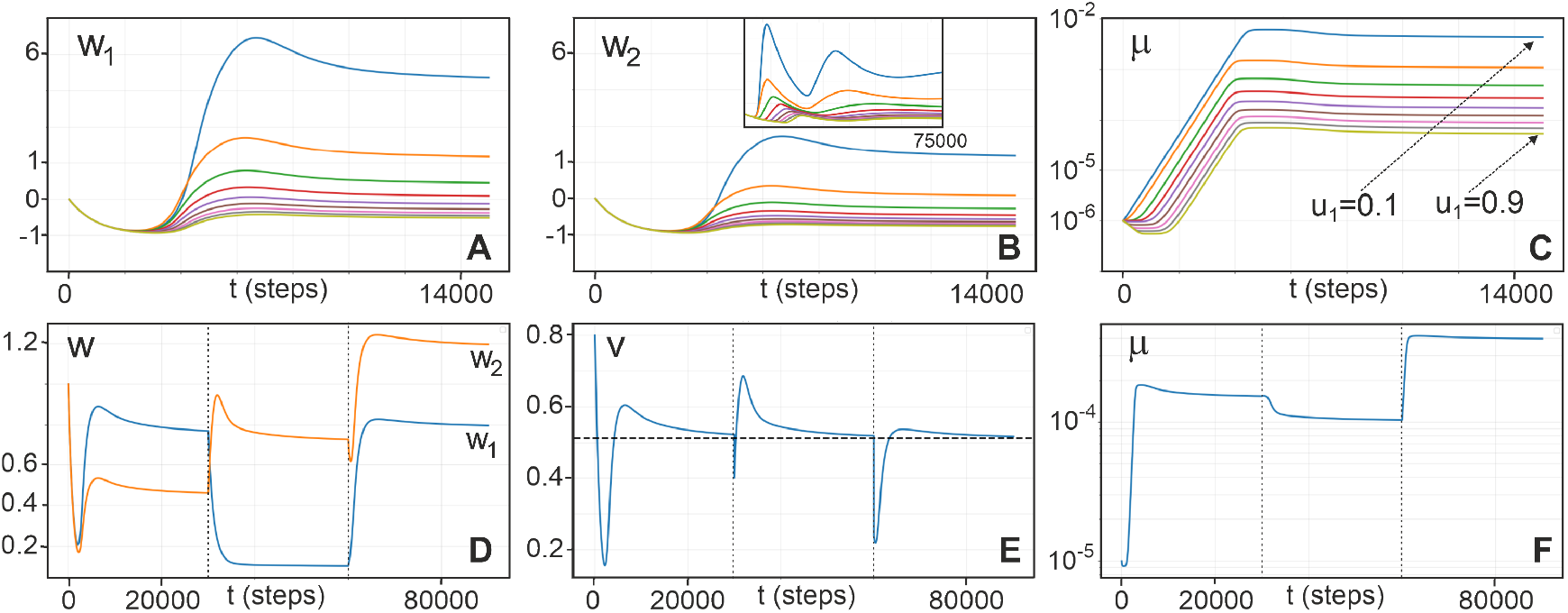
Temporal development of this system calculated using Runge-Kutta integration with a step of 0.2. For parameters see Eq. 38. Initial conditions *w*_1,2_(0) = 1.0 and for panels (A-C): *µ*(0) = 10^−6^ while for (D-E) *µ*(0) = 10^−5^. For A-C we used *u*_1_ ∈ [0.1, 0.9] in steps of 0.1 as indicated in (C). The inset in (B) shows the behavior of *w*_2_ for *τ*_*w*_ = 1.0. Note the much longer time axis here. Panels D-E show the behavior when suddenly changing the input situation. We have used *u*_1_ = 0.5, *u*_2_ = 0.3 for *t* = [0, 30000]; *u*_1_ = 0.1, *u*_2_ = 0.7 for *t* = [30000, 60000] and *u*_1_ = 0.2, *u*_2_ = 0.3 for *t* = [60000, 90000]. The dashed line in (E) is at *v*^∗^ = 0.514. Note that the same fix point exists for panels A-C (not shown).

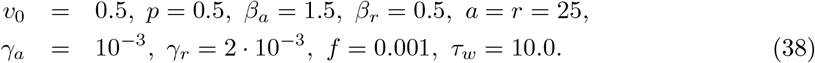

For this we find that *v*^∗^ = 0.514 (Eq. 46) and obtain the curves in Figure 3 for the other three fix points depending on *u*_1_.

Figure 4 shows the time-development of this system. Panels A-C demonstrate that convergence is obtained for different constant inputs *u*_1,2_ after about 10.000 time steps. Panels E-F show the relaxation behavior when suddenly changing the inputs. Convergence takes a bit longer now until about *t* = 20.000. In (E) it can be seen that the system in all cases approaches the same stable fix point *v*^∗^ = 0.514 (dashed line).

Figure 5 demonstrates that this system can also follow fast changing inputs (A-C). Here we used constant *u* only for 250 time steps each. After this, the input could change where three situations with equal probability had been used: (*u*_1_, *u*_2_) = (0.3, 0.0) or (0.0, 0.4) or (0.3, 0.4). The inset in (C) uses the sum of *u*_1_ + *u*_2_ to show how this looks like up to time step 3.000. The average activity (B) matches the expected fix point (here *v*^∗^ = 0.588). Panels (D) and (E) represent the distributions of weights *w*_1,2_ over time and interestingly in panel (F) we show that there remain three distinguishable clusters of activity *v*, where the leftmost corresponds to (*u*_1_, *u*_2_) = (0.3, 0.0), middle to (0.0, 0.4) and right to (0.3, 0.4). Hence, the system can discriminate these situations “sorting” the outputs according to input strengths. We note that this “sorting property” has been a major advantage of the original ALL rule [12]. Different from the other rules to which it had been compared it creates – and here this holds, too – an ecologically useful ordered output structure.

**Figure 5.**
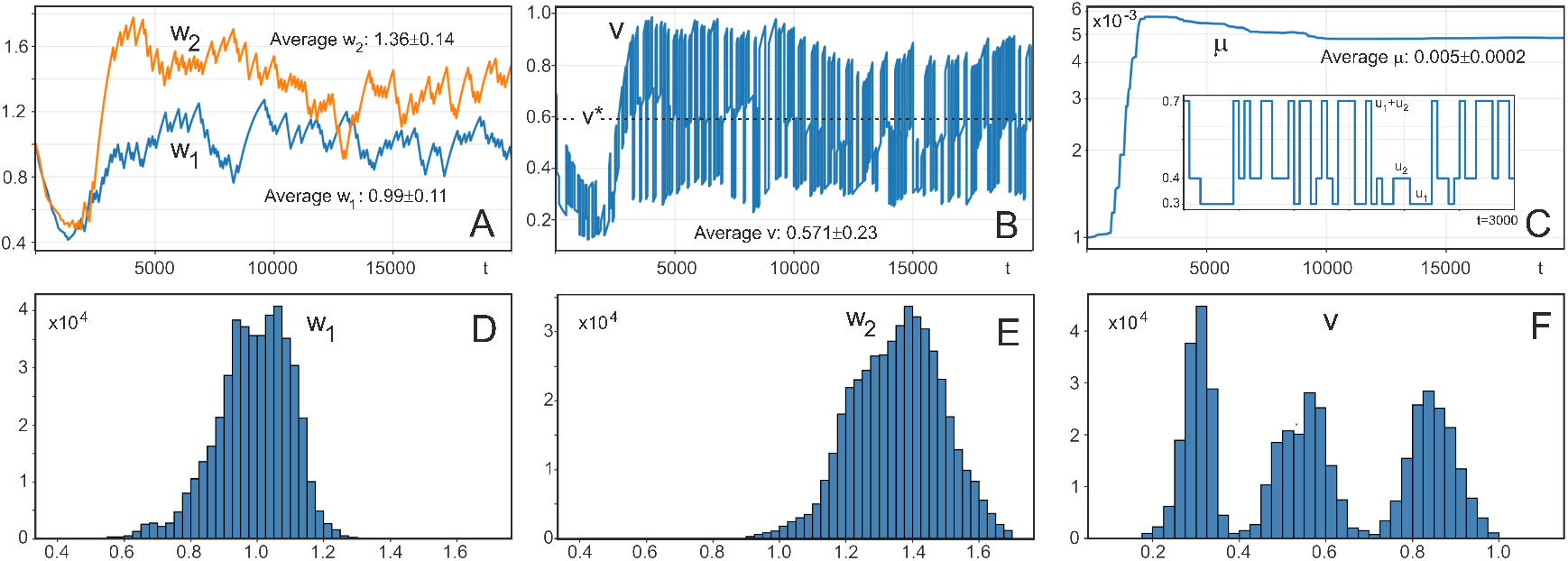
Temporal development for quickly changing inputs with a total simulation duration of *t* = 100.000. Inputs *u*_1,2_ were changed every *t* = 250 steps equally distributed with probability 1/3 each as: (*u*_1_, *u*_2_) = (0.3, 0.0) or (0.0, 0.4) or (0.3, 0.4). The inset in panel C shows this exemplary using their sum for up to *t* = 3.000. Panels A-C show the time courses up to *t* = 20.000, D-F show histograms calculated for the period *t* ∈ [10.000, 100.000] where the system has basically converged. Averages were also calculated for the same period. A) weight development, B,C) development of *v* and *µ*. D,E) histograms for the weights and (F) for *v*.

Note that convergence time of weights and output is not much affected by the number of inputs. Figure 6 shows two pairs of curves each with *I* = 20 inputs of different strength and with different input probabilities *p* (see legend for details). The system is essentially converged after about 5000 steps. Panels A,B show a case where 4 inputs had a higher *p* than the rest. The weight plot (B) compares each higher-p-input (4, 9, 12, 15) with its “next neighbor” (5, 10, 13, 16). Next neighbors have a slightly higher input *strength* (see legend) but nonetheless higher-p-inputs reach - much as expected - larger weights. For example: Compare the red curve for *w*_15_ with the gray one for *w*_16_. Also here we find that convergence times remain the same. The table on the right side shows that after 10000 steps the average of *v* reaches the expected value of *v*^∗^ = 0.588 in all cases within statistical limits. Convergence times will also be assessed below for the navigation task and in the Discussion we will compare this to real time.

**Figure 6.**
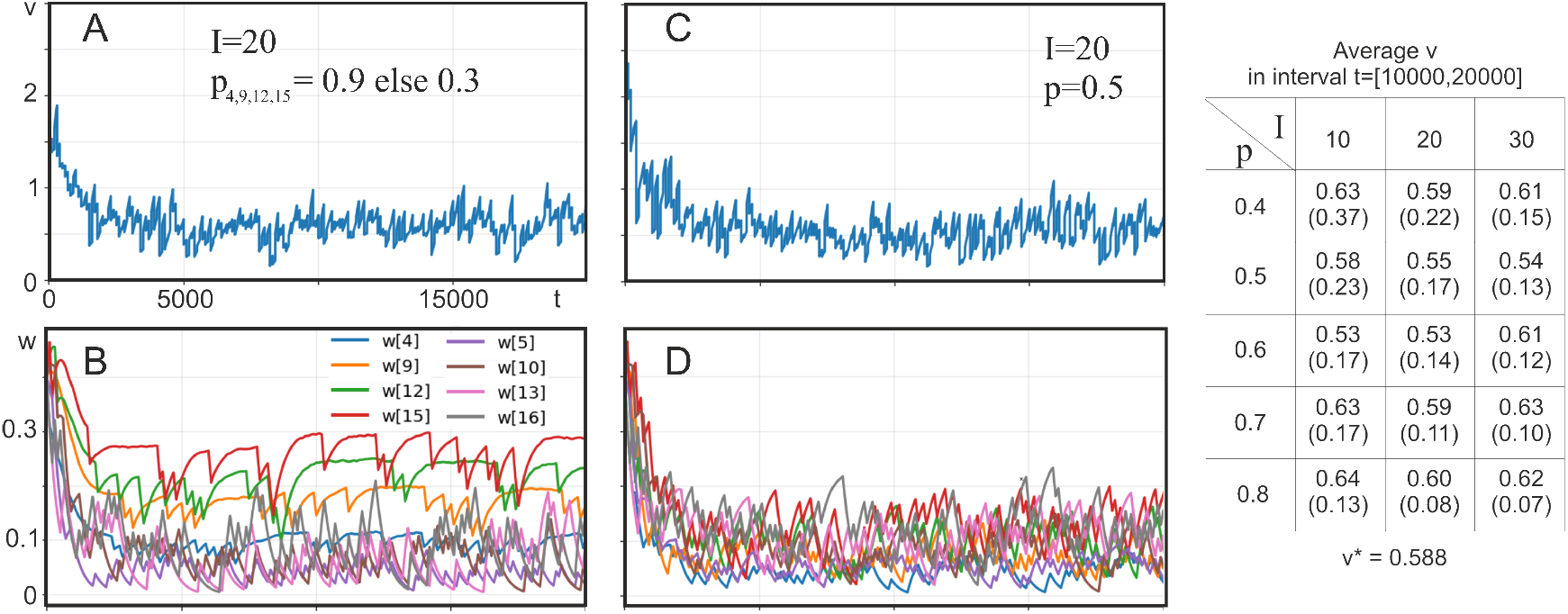
Convergence characteristics for many variable inputs. Total simulation duration of *t* = 60.000. Panels A-D show outputs *v* and 8 selected weights *w* for *I* = 20 inputs. The table summarizes average values for *v* for *I* = 10, 20, 30, with standard deviations in parentheses. Procedure: In general inputs were either constant with a certain value or zero, where the probability *p* for taking a non-zero value was set by us. In panels A,B non-zero inputs 4, 9, 12, 15 had a probability of *p* = 0.9 all others *p* = 0.3, in panels C,D and for the table all probabilities where *p* = 0.5. The actual numerical values for non-zero inputs had been determined by dividing the full input interval *U* = [0.1, 0.9] into *I* non-overlapping, small sub-intervals *U*_*i*_ with *i* = [1, …, *I*], where the first starts at 0.1 and the last ends at 0.9. Every input *u*_*i*_ had then been associated to its sub-interval *U*_*i*_. Hence, the first input *u*_1_ gets values from within 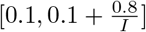 and the last *u*_*I*_ from within 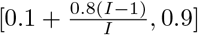. Following this assignment the actual input values had been chosen randomly with these sub-intervals. For 100 time steps, the belonging input was then set to this value with a chance of *p* or to zero, else. Parameters as before: *a* = *r* = 45.0, *γ*_*a*_ = 10^−3^, *γ*_*r*_ = 10^−2^, *β*_*a*_ = 2.0, *β*_*r*_ = 0.25, *v*_0_ = 0.5, *τ*_*w*_ = 2.5, *f* = 10^−3^. Initial conditions were *w*_1,2_(0) = 0.4 and *µ*(0) = 10^−3^. Note that *v*^∗^ = 0.588 as before.

## Navigation using learned place fields

The property of forgetting coupled with learning rate recovery should allow an agent to adapt efficiently to a changing environment. To test this we had designed a system (Fig. 2) were an artificial agent learns locating a food source using odor as well as location inputs (via place cells). The goal was to allow the agent to learn a new food location after having targeted (and “consumed”) the currently existing one. To this end the agent first explores the arena with a direction-preserving random walk (Fig. 7 A) and generates a place field system at the level of the 2nd-order place cells (see top of Fig. 2) containing an activation-gradient with the strongest activations found near the food source from which on they drop with distance (Fig. 7 B). After some time the agent can switch to exploitation and target the food source directly, where we show paths from three different, randomly selected starting points to the food source overlaid onto the place field system in panel B, too. To compute these paths, the agent follows the activation gradient as explained in the subsection “Exploitation Mode”, above. Note that here and in the following we show results using a normalized arena size of 1 × 1 and also (next) a time axis that uses simulation steps. Please see the Discussion section for an estimate how this would map to real time and space.

**Figure 7.**
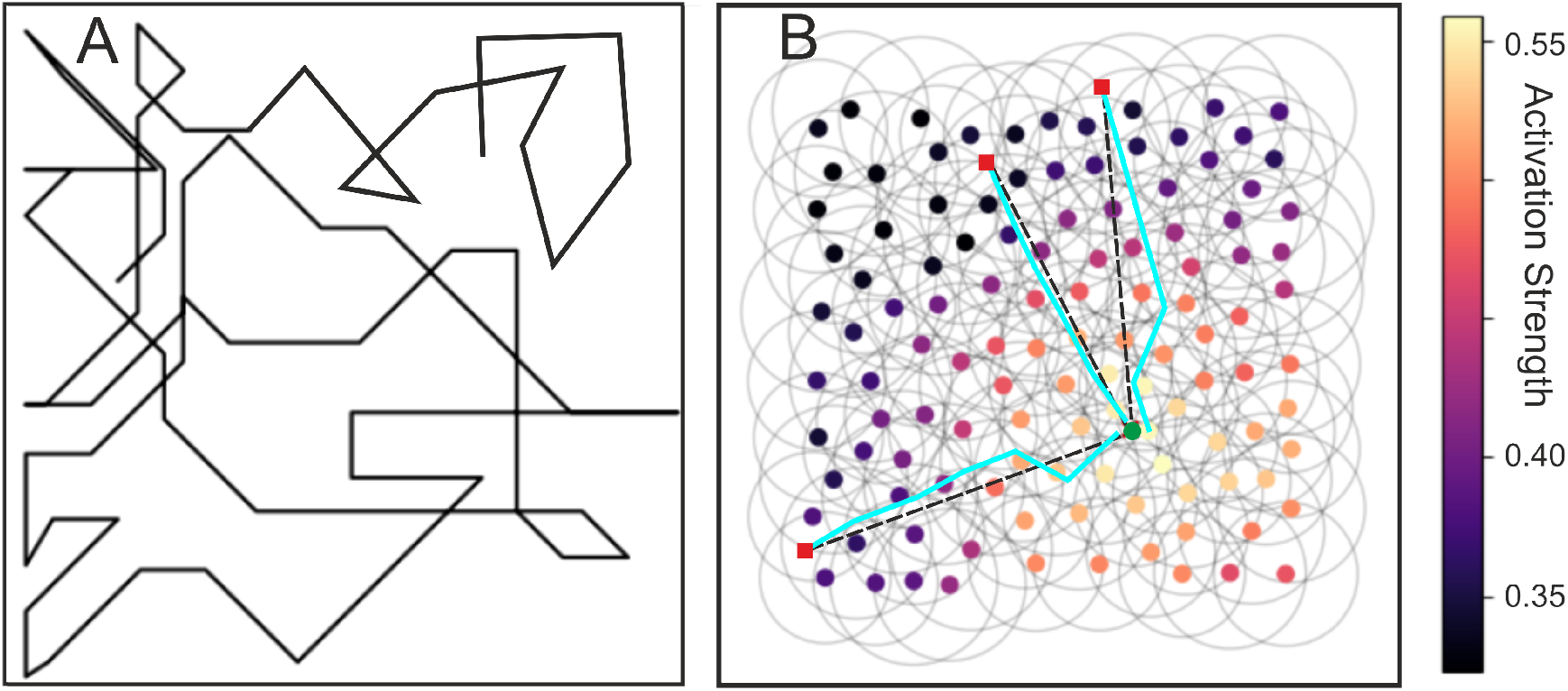
One path that shows an exploration pattern (A); and (B) the resulting layer 2 activations leading to a gradient from repeated exploration towards a food source (green) where 3 paths during exploitation are superimposed from different starting points (red). Black dashed straight lines represent the shorted possible paths. For this and for figures 8,9 we have used these parameters: *a* = 50, *r* = 50, *γ*_*a*_ = 10^−1^, *γ*_*r*_ = 10^−2^, *β*_*a*_ = 0.8, *β*_*r*_ = 0.25, *τ*_*w*_ = 1, *f* = 10^−2^, *v*_0_ = 1.0.

The above results are based on the characteristics of the layer 1 place cells that converge onto the ones in layer 2. Hence, the belonging temporal development of weights and activations of the different cells during the exploration phases is shown in Figure 8 A-G. The food source had been twice relocated at *t* = 15.000 and 30.000. (Note that exploitation only needs single runs and would follow at the end of exploration, see Fig. 7 above.) The inset in panel (I) shows the locations of the food source (stars) and 10 selected place field center locations as colored dots. Color coding in all panels correspond to these place fields. For example, the top row (A-C) shows weights and activations of 10 selected units from the stack of 200 cells in layer 1, which belong to the red place field. As expected, when the place field is close to the odor source, which is the case for source location 2, then the odor weight will increase. Due to the fix point characteristic of our rule the place field weights will then drop to some degree such that the activations in layer 1 will indeed remain close to the expected fix point, which can be calculated using the solution from above as *v*^∗^ = 0.502. In panels (D) and (E), this antagonistic behavior of place field- and odor-weights can be seen for all locations from the inset and corresponds to the behavior found for two inputs in Fig. 4 D. Learning rates in layer 1 (panel F) drift to some degree in accordance to the noisy response patterns of the layer 1 units.

**Figure 8.**
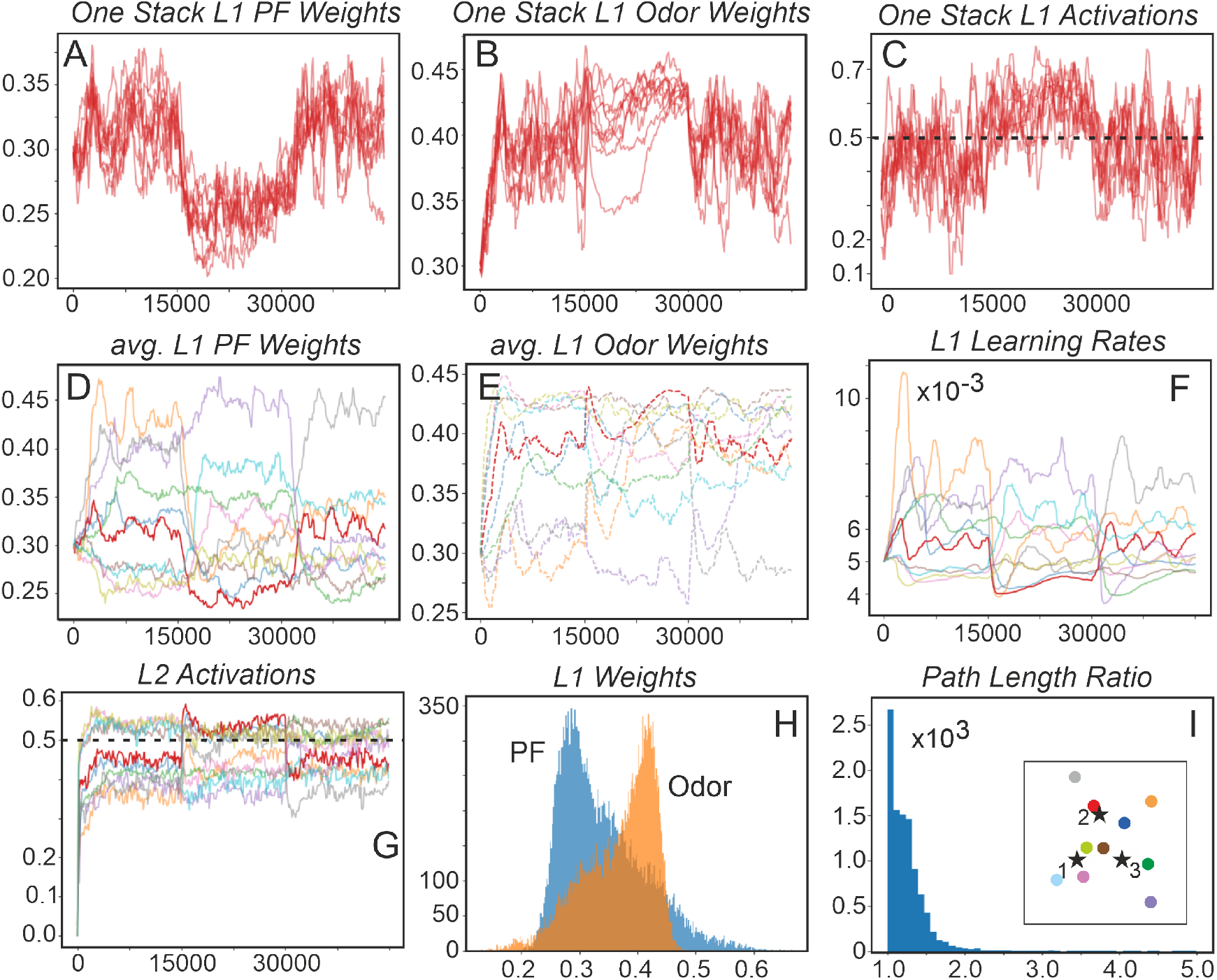
Weight and activation development when relocating the food source at *t* = 15.000 and 30.000. The inset in panel (I) marks with a star three food source locations and with colored dots those 10 locations for which we show the weight and learning rate development in the different panels. A-F) Time-courses of weights, activations and learning rates in layer 1. Panels A-C show 10 randomly selected cells from the layer 1 stack that belongs to the red place field location (see inset in I). The dashed lines in panels (C) and (G) represent the analytically calculated fix point *v*^∗^. D-F) Results for all 10 locations. In (G) we show the layer 2 activations and H,I) show histograms of weights after the exploration phase and path-length ratio for 10000 individual weight development for ten place cells in layer 1.

**Figure 9.**
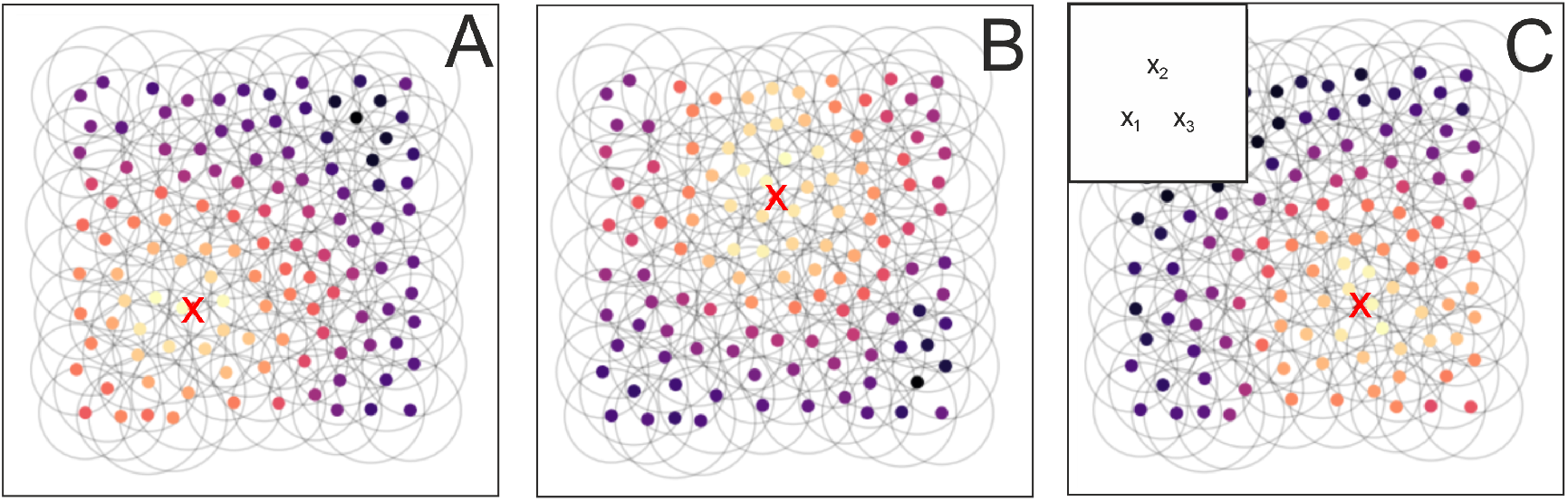
Relocating the food source twice (at *t* = 15000 and *t* = 30000) and relearning the 2nd-order place-cell activations. Color scale as in Figure 7. The inset in (C) marks with “x” three food source locations. A) Starting situation, where the red cross marks food source *x*_1_, B) first relocation to *x*_2_, C) second relocation to *x*_3_.

Averaging nearby stacks (and low pass filtering) creates the receptive fields of the layer 2 place cells with their activity. Here we note that layer 2 activity (Fig. 8, panel G) still relates to layer 1 (and, thus, remains within reasonable limits), but it begins to loose the fix point characteristic, which is important as the system needs to represent with its activity the proximity of a place field to the food source. We have consistently observed that layer 2 converges onto activity levels, which correlate to the distance of a place field to a food source leading to a gradient and, thus, to convergent paths during exploitation (see statistics in Fig. 10). This can be explained by the property that layer 2 units perform activity sampling from the stack(s). Note that units in each layer 1 stack only respond with 0 or 1 times their weights (see Eq. 16). Hence, whenever the units in a stack are close to a food source, then on average more cells in this stack will for every time step produce a non-zero response leading to a higher response in layer 2 as compared to a situation when the food source is far away. This behavior can be clearly seen, for example for the red cell in Fig. 8, panel F, where the 1st and 3rd food sources are far away from this place field, where the 2nd is nearby. Hence, while layer 1 will roughly follow the fix point characteristic, with sampling we are still able to arrive at location-specific responses in layer 2. To control for this we had performed an experiment where we had used only one layer with continuous-valued place cell characteristics (using bell-shaped kernels). Here cells will indeed again follow the fix point characteristic but such a system cannot produce any location-specificity (data not shown).

**Figure 10.**
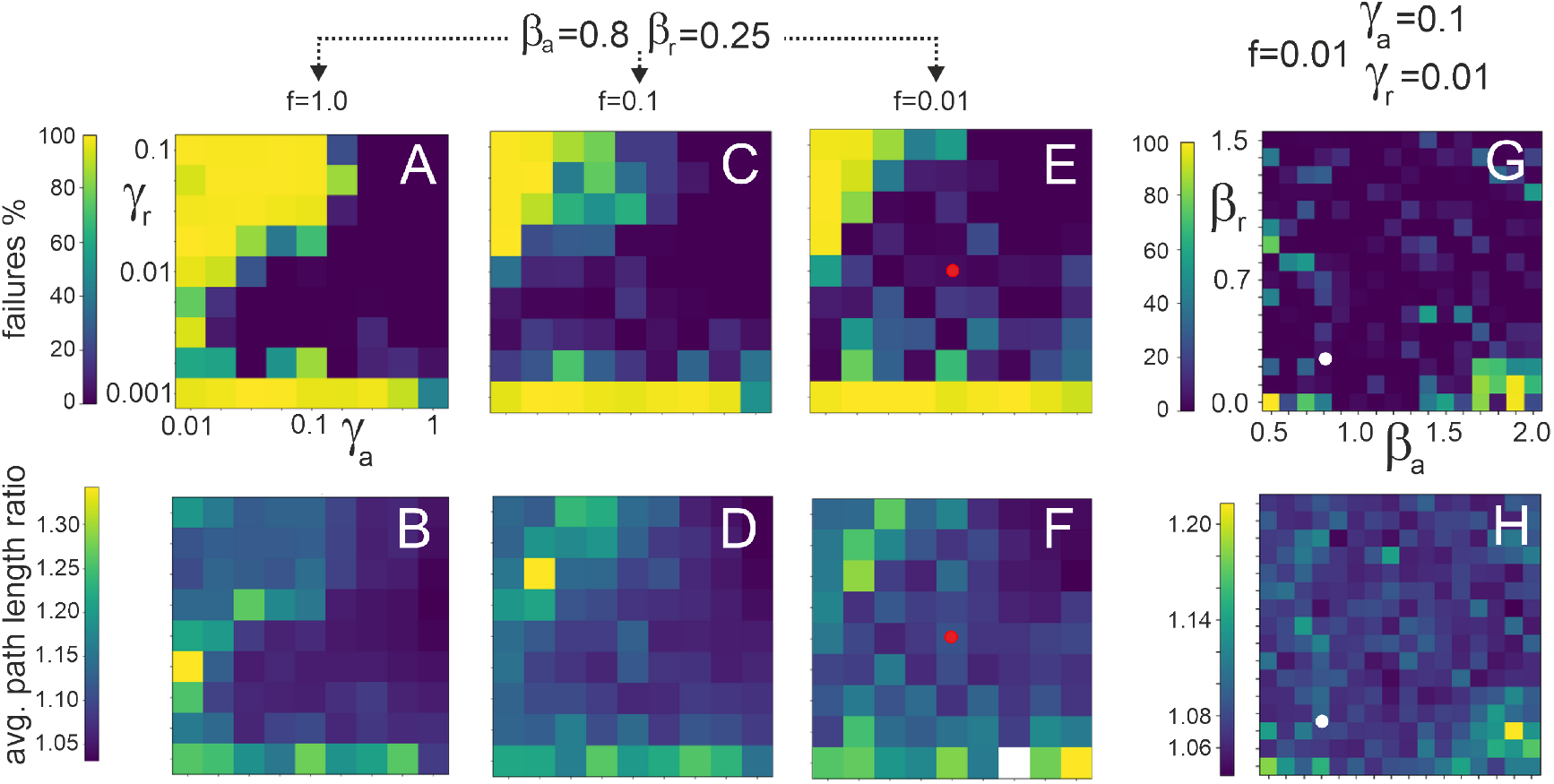
Statistical evaluation of different parameters. The top row shows in percent how often paths could not be found, the bottom row the average path length ratio relative to the optimal (straight) path. Parameters as indicated at the panels, all other parameters were as before. The red dots marks the parameters used for panels G,H, the white dots those used for (A-F). The white pixel in panel F indicates that here no converging paths had existed at all (100% failures).

Panel (H) shows in summary the antagonism of place field-versus odor-weights in layer 1, as discussed above, for all weights obtained during this run. The path-lengths of the exploitation phase remain short with a mean ratio *R* of actual-divided by optimal-path length of 1.55 (panel I), but a few very long paths exist because the agent sometimes oscillates between two place fields when the gradient is not steep enough.

In Figure 9 we show three different 2nd-order place field systems with their activation patterns at the end of the exploration phases, resulting from relocating the food from location *x*_1_ to *x*_2_ and then to *x*_3_.

Figure 10 shows statistical results on parameter sensitivity asking about convergence as well as path lengths in our 1.0 × 1.0 arena. For this we have evaluated 10, 000 paths for every parameter combination. Minimal allowed path length was 0.3 to exclude nonsensical mini-paths from the statistics. Furthermore, starting points of the agent had to be away from each arena border by 0.1 to make sure that every start point would hit at least one place field. One finds (top row) that within rather large ranges the system converges robustly. For *γ*_*r*_ = 0.001 the system will not converge any longer as can be seen by the yellow line at the bottom of each plot (A,C,E). Here recovery is too weak and the gradient across the place fields in the arena becomes diffuse. Conversely, we also observe that the system fails when recovery is similar to or stronger than annealing leading to the yellow triangle in the top left of these plots. In these cases the too-strong recovery leads to divergence of the activity levels in the system. Note that even for these parameter settings there were still several converging paths (hence 100% failures did in fact not occur) on which the path-length analysis in the bottom row (B,D,F) is based (only for one case we had 100% failures, see white pixel in panel F). In general the bottom row shows that, when the system converges, then the resulting paths remain short. At the right side we investigated the influence of *β* and found that this has little influence over a wide range, too.

## Discussion

In this study we have introduced an unsupervised synaptic plasticity rule that uses several ecologically justifiable aspects. For this we have combined forgetting of synaptic weights *w* with a process of meta-learning, which does not affect weights directly but rather their change-rate *µ*. Here we assume that the change-rate should go up, allowing weights to grow, when the neuron’s output is too small and vice versa in case the output gets overly strong. The system introduced here also makes use of a modified learning rule, where we have reduced the influence of the neuronal output onto learning to an all-or-none behavior by using the Heaviside function. This way learning starts as soon as the neuronal activity is larger than some threshold *v*_Θ_ (here we had kept this, for convenience, at zero) but does not depend on the actual magnitude of the neurons output. As a consequence weight growth is now linear and not exponential as in the conventional Hebb-rule (see [12] for a more detailed analysis of this).

There are two main findings to consider:

1. We have shown that the neuron’s output remains, after learning, close to a predetermined (“target”) value and this is achieved independent of the input(s). This property can be understood as “synaptic scaling” [8]. Experimental evidence supports that neurons ‘want” to achieve a certain target activity [8, 10] and that synaptic changes are driven towards achieving this target. Our system does this and approaches *v*^∗^ independent on the input as a stable fix point. Noteworthy, too, is the fact that convergence is quite fast and independent of the number of inputs (Fig. 6) leading to the fact that the system can adjust multiple synapses (e.g., in the simulated place cell network) fast enough to allow for the learning of different environments.
2. In addition to this, the system keeps outputs sorted according to their input strengths (see Fig. 5 F), following the properties of the original ALL rule [12]). Intuitively, this seems to be a straight-forward requirement for any learning system: Stronger inputs (or stronger conjoint input combinations) should lead to stronger outputs. Remarkably, this is however, not the generic case and other – often used – learning rules (like BCM, Oja’ rule, conventional scaling rule) cannot achieve this (for analysis see [12]).

Finally we had applied our learning method to a simulated food-targeting task using a rather strongly abstracted version of a place-cell system. Here we were not interested in true physiological realism and the place cell system we used does not very deeply reflect current knowledge. Instead, we were concerned with the question whether or not our meta-learning system would still produce useful results in a bigger network with changing environment.

### All-or-none learning with a forgetting term (Eqs. 41, 42)

Using a Heaviside function for Hebbian learning has clear theoretical advantages, as it yields only linear weight growth. In contrast, the conventional Hebb rule, driven by membrane potential, produces exponential growth and runaway strengthening of dominant inputs. Note that post-synaptic depolarization—especially at dendritic spines—affects *Ca*^++^ influx through NMDA channels in an all-or-none manner that governs plasticity. Estimated thresholds are 150–500 nM for LTD and *>* 500 nM for LTP [16]. A single EPSP can raise *Ca*^++^ to about 700 nM, while pairing with post-synaptic depolarization can reach up to 12 *µ*M [17]. All this is compatible with an all-or-none process (at least at spines).

Models of spine plasticity predict that even a single synaptic EPSP can trigger significant NMDA-mediated *Ca*^++^ influx [20], consistent with experimental findings [21–23]. Thus, back-propagating or dendritic spikes are likely sufficient to induce plasticity at a spine. This supports a sharp transition in post-synaptic learning effects, with the Heaviside function as a limiting case. Steep sigmoidal functions could be used instead, with minimal impact on results.

The forgetting term that appears in our learning rule is nothing special as it reflects the generally known aspect that synaptic weights will decay when relevant inputs are missing. This notion is related to the effect of “extinction” and can be found already in the work of Pawlov as well as Hull [24, 25] and in the first models for this by Rescorla and Wagner in 1972 [26]. **Meta-Plasticity in the form of learning rate annealing (reduction of the learning rate) and recovery (Eq. 43):** Meta-plasticity had been discussed starting mainly in the 1980th (e.g. *depotentiation* with [27] and without [28] LTP). See [29, 30] for early reviews of this and other meta-plasticity aspects. Several processes support “plasticity of plasticity”. For example. in 1998, Bi and Poo [15] showed that changes in EPSP amplitude are inversely related to its initial size, so larger synapses grow less than smaller ones. This likely reflects an LTP saturation (ceiling) effect and could, in theory, be modeled as learning-rate annealing. However, this exposes a key issue: for theorists, the learning rate is a single abstract variable and annealing+recovery represents a simplification of complex meta-plasticity, making links to underlying biophysics difficult. Many studies suggest that LTP reduction but also facilitation may involve NMDA receptor–related mechanisms [31–35], but these effects may be too short-lived, decaying within about an hour [31]. Stimulus-driven (dis-)facilitation of LTP, however, also operates over longer timescales [36, 37] covering effects for up to seven hours. More persistent LTP reduction may also arise from mechanisms governing late-LTP [38–41], which are thought to underlie synaptic consolidation. Yet their role in meta-plasticity remains unclear. Still, it is plausible that neurons reduce “learning effort” by down-regulating key biochemical components via saturation-like kinetics once activity is sufficiently high, effectively implementing learning-rate reduction. Hence in conclusion, effects that can be interpreted as learning rate reduction as well as -recovery are indeed abundant and the here-introduced equation for this can be understood as an abstraction of such processes.

Considering theoretical approaches, simulated annealing has long been established as a standard technique in reinforcement learning (RL), where it is commonly employed for purposes such as adaptive step-size reduction [42], the gradual decay of exploration rates [43], and exploration scheduling in deep-RL frameworks [44]. Beyond RL, annealing approaches are also extensively used in supervised learning settings [45–47], as well as across several variants of Hebbian learning [48–51]. The latter line of work is particularly relevant to the present study due to its focus on self-organized adaptation mechanisms. In most of these previous studies, however, annealing is introduced primarily as an auxiliary mechanism that facilitates stable and efficient convergence of synaptic weights or network parameters. By contrast, we are not aware of prior work that systematically investigates annealing in conjunction with other methods as the key mechanisms underlying the stabilization of neural activity dynamics.

### Responses to a changing environment

Part of this study was devoted to use this learning framework in a bigger network, which was inspired by the hippocampal place cell system. Already early-on the place field system has been considered and modeled as a map for navigation [52] and an influential model based on attractor neural network had suggested a mechanism for this [53]. The combination of place field and odor information had also been modeled some time later [54].

While these and other contemporary models [55] pay much more attention to physiological details as compared to the network used here, our approach was focused on the question whether or not the theoretical properties that we had observed would still hold here, too. Our results demonstrate this and we found that the system is quite robust against parameter variations.

It is of interest to estimate how the here-observed behavior in simulated space and time would map to reality. We assume an arena size of 1×1 m and place field diameters of 15 to 30 cm, which holds for rats in small environments [56, 57]. We have set the speed of the agent to 10 cm per simulation time step and we assume a desired average movement speed of 40 cm per second. This maps to 4 simulation time steps per real second. One epoch for presenting a specific food source is 15,000 time steps corresponding to 3750 seconds of total exploration time. From Figure 8 one can estimate that curves become reasonably flat (converged) after about 3500-4000 time steps (see panel G), where this takes longer for the first epoch due to the required initialization of the system. Hence time to convergence would be about 875-1000 seconds and the traveled distance of the agent less than 200 m. Note that this leads to a *full* exploration of the arena. In reality animals will, very likely, stop exploring as soon as a “good” path from their burrow to a food source exist. Hence, taking this into account, too, the here-calculated mapping between the simulation of a full exploration and reality appears very reasonable.

In summary, the here-introduced learning rule combines several physiologically grounded mechanisms. It results in stable input-independent fix points, which represent the property of synaptic scaling. Furthermore, inputs and their combinations lead to outputs which reflect the input strengths. The fact that learning is balanced with forgetting allows also for quite fast adaptation to a changing environment. The rule contains indeed quite a few parameters but it was found to be robust within wide ranges.

## Acknowledgments

The authors acknowledge financial support from the European Union’s Horizon Europe programme through the FlexCycle Project (Grant Agreement No. 101189600). We are grateful for discussions with Hannah Kerger on the topic of forgetting and to Lennard Jahn for discussing the relevance of different parameter ranges with us.

## Appendix

### About the limit towards infinity of variable a

Co nsidering: 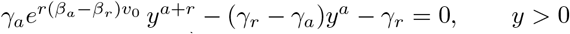. Factor out 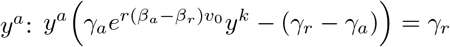. As *a* → ∞, if *y <* 1 then *y*^*a*^ → 0, and if *y >* 1 then *y*^*a*^ → ∞; in both cases the left-hand side cannot balance the finite constant *γ*_*r*_. Hence a finite nonzero limit requires *y* → 1. Therefore lim_*a*→∞_ *y* = 1. and, thus, lim_*a*→∞_ *v*^∗^ = *β*_*a*_*v*_0_.

### Stability of the non-trivial fix point

Most straight-forward condition: An FP is stable if all Eigenvalues of the Jacobian of the FP are smaller than zero. We consider the system given by:

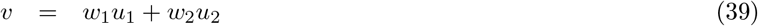

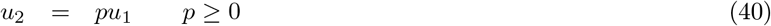

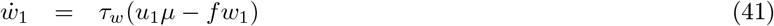

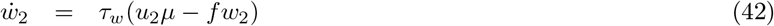

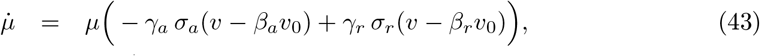

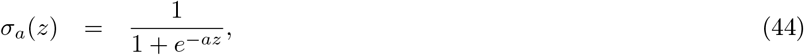

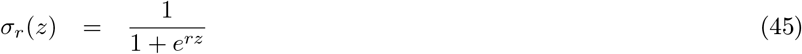

With fix points:

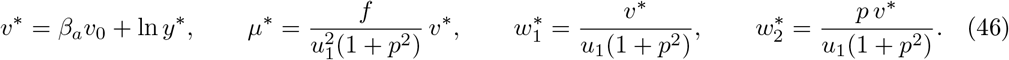

and from Eq. 43 we have:

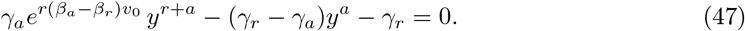

Nontrivial fixed point *µ*^∗^≠ 0, thus renders:

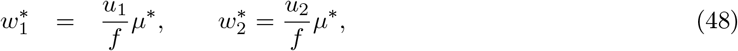

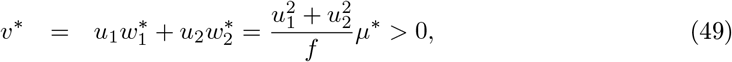

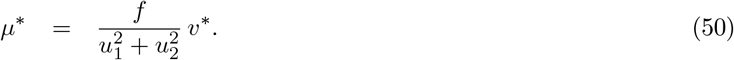

**Jacobian:**

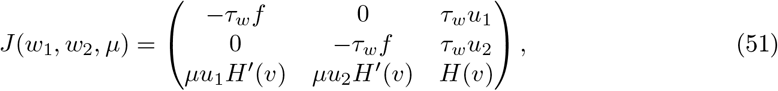

with:

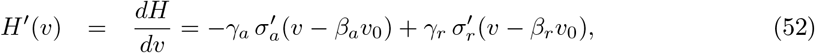

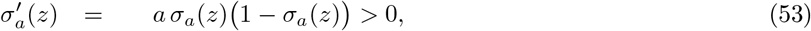

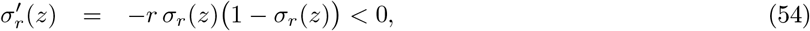

Leading to

⇒ *H*^′^(*v*) = *γ*_*a*_*a σ*_*a*_(*v β*_*a*_*v*_0_) 1 *σ*_*a*_(*v β*_*a*_*v*_0_) *γ*_*r*_*r σ*_*r*_(*v β*_*r*_*v*_0_) 1 *σ*_*r*_(*v β*_*r*_*v*_0_) *<* 0.

Thus, in short (needed at the bottom):

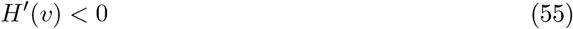

**At the nontrivial fixed point:** *H*(*v*^∗^) = 0

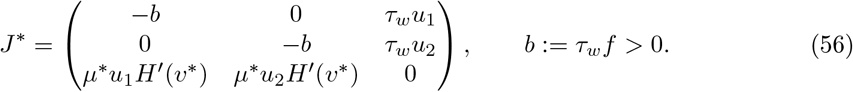

The characteristic polynomial is

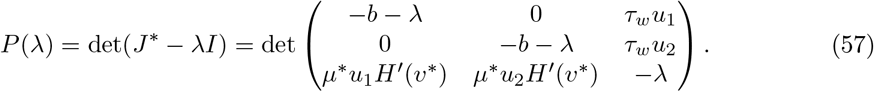

This can be calculated step-wise and results in the following Eigenvalues:

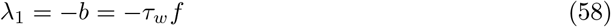

where *λ*_2,3_ are the roots of the quadratic

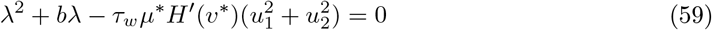

Thus:

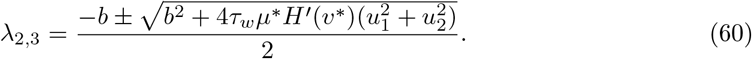

Using the fixed point identity 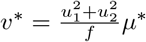 (so 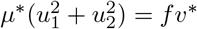), this can also be written as

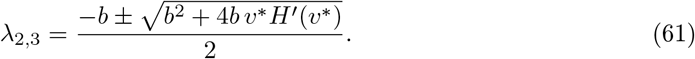

Under the above conditions *τ*_*w*_ *>* 0, *f >* 0, *b* = *τ*_*w*_*f >* 0 we find that 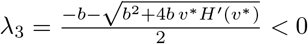 always and

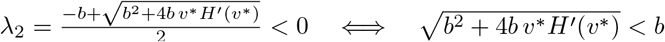.

Squaring: 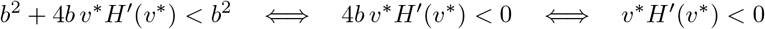.

Hence:

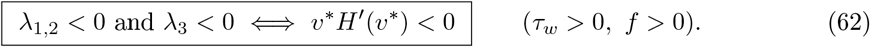

We know that *v*^∗^ *>* 0 and form Eq. 55 that *H*^′^ *<* 0. Hence all Eigenvalues are negative and the fix point is stable.

